# SPREd: A simulation-supervised neural network tool for gene regulatory network reconstruction

**DOI:** 10.1101/2023.11.09.566399

**Authors:** Zijun Wu, Saurabh Sinha

## Abstract

Reconstruction of gene regulatory networks (GRNs) from expression data is a significant open problem. Common approaches train a machine learning (ML) model to predict a gene’s expression using transcription factors’ (TFs’) expression as features and designate important features/TFs as regulators of the gene. Here, we present an entirely different paradigm, where GRN edges are directly predicted by the ML model. The new approach, named “SPREd” is a simulation-supervised neural network for GRN inference. Its inputs comprise expression relationships (e.g., correlation, mutual information) between the target gene and each TF and between pairs of TFs. The output includes binary labels indicating whether each TF regulates the target gene. We train the neural network model using synthetic expression data generated by a biophysics-inspired simulation model that incorporates linear as well as non-linear TF-gene relationships and diverse GRN configurations. We show SPREd to outperform state-of-the-art GRN reconstruction tools GENIE3, ENNET, PORTIA and TIGRESS on synthetic datasets with high co-expression among TFs, similar to that seen in real data. A key advantage of the new approach is its robustness to relatively small numbers of conditions (columns) in the expression matrix, which is a common problem faced by existing methods. Finally, we evaluate SPREd on real data sets in yeast that represent gold standard benchmarks of GRN reconstruction and show it to perform significantly better than or comparably to existing methods. In addition to its high accuracy and speed, SPREd marks a first step towards incorporating biophysics principles of gene regulation into ML-based approaches to GRN reconstruction.

## INTRODUCTION

Gene regulatory networks (GRNs) are a popular framework for describing mechanisms underlying transcriptomic changes associated with a variety of biological processes, such as development [1, 2], behavior [3] and cancer [4]. GRNs catalog transcription factors (TFs) and their target genes, each such TF-gene relationship represented as an edge in a network. Typically, such an edge means that the TF directly regulates the gene, i.e., perturbing the TF’s concentration should change the gene’s expression and that such a causal relationship arises from the TF binding to an enhancer or promoter associated with the gene. Accordingly, methods for reconstructing GRNs rely on statistical relationships between TF and gene expression, as well as evidence of TF-DNA binding that might underlie those relationships [5, 6]. However, it is often the case that the researcher only has access to gene expression data for their system, with other types of data such as TF-DNA binding profiles, histone modification profiles, and 3D chromatin data being unavailable [7]. Thus, GRN reconstruction solely from expression data is an important open problem, and the focus of our study.

Given a matrix of expression values, with rows representing genes (including TF genes) and columns representing different conditions or cells that have been profiled with transcriptomics technologies [8], the task is to infer the underlying GRN. This amounts to detecting covariation between expression levels of TF and gene (rows) across the different measurements (columns), and the biological interpretation of an edge is thus contingent on the kind of variation represented by columns of the matrix. The nature of variation and covariation present in expression matrices differs substantially between single-cell and “bulk” transcriptomic datasets, leading to GRN inference methods specialized for either domain. For instance, methods for the single-cell domain must tackle high levels of technical noise [9] but have very large numbers of samples (∼1000 to ∼100,000 cells) to rely on, while GRN inference from bulk transcriptomic data must work with few samples (∼10 to ∼100 conditions or individuals) but do not face the challenge of dropouts [10]. Here, we focus on GRN inference for bulk transcriptomics data, though the core ideas are germane to the single-cell domain as well.

Many statistical and machine learning approaches have been proposed for GRN reconstruction from expression data, including those based on correlations and information theoretic measures [11–13], probabilistic graphical models including Bayesian networks [14–18], Boolean networks [19, 20], differential equations [21], linear regression [22–24], Random Forests [25, 26], gradient boosting [5, 27, 28], neural networks [29–34] among others. The more popular approaches are based on the idea of training a multi-variable model that predicts a gene’s expression from the levels of TFs, reporting the TFs that are most useful for such prediction as the regulators of the gene (A separate model is trained for each gene). A major challenge faced by these methods is that the number of covariates is typically far greater than the number of samples, making it difficult to train the models robustly [35]. A second, related limitation of the approach is that the mathematical “form” of the relationship between TFs’ and gene expression must be pre-determined and encoded into the structure of the model, e.g., as a weighted sum, decision tree, etc., and rich representations that accommodate a greater variety of possible relationships, e.g., neural networks, must be eschewed due to small sample size.

Recent studies have explored a different line of attack on the problem: that of training a model to directly predict a TF-gene relationship from expression profiles of a gene and its candidate regulator. The key difference here is that a “sample” (unit for which a prediction is to be made) is now a TF-gene pair rather than a biological condition, and the output of the model is the presence or absence of regulatory relationship between that TF-gene pair, rather than gene expression in a condition. This addresses the small sample size of above-mentioned methods, since the number of TF-gene pairs is large, but poses a different challenge: that of determining a training set where many TF-gene pairs are labeled as having true regulatory relationships or not. One possible solution, adopted by Yuan & Bar-Joseph [29], is to use epigenomic evidence of TF-gene relationships (e.g., TF ChIP-seq) to define positive examples for training. With a large enough training set of TF-gene pairs and their positive/negative labels, various classes of ML models, including deep neural networks [29, 33], may then be trained to predict GRN edges from the joint distribution of TF and gene expression. A distinct advantage of this approach is that it offers greater flexibility in modeling the relationship between TF and target expression. In this study, we sought to further explore this emerging approach to GRN inference, which departs significantly from the dominant paradigm today. We will refer to this approach as “supervised GRN reconstruction” since it relies on examples of true and false GRN edges to train predictive models.

Despite the promise of its early versions [29], a major limitation of the supervised GRN reconstruction strategy is in setting up the training set. This is because “gold standard” (highly accurate and comprehensive) GRNs are almost non-existent today. Indeed, the most widely adopted strategy for benchmarking GRN inference tools, including in community-wide efforts such as DREAM challenges [36], continues to rely on synthetic data sets. “Real data” benchmarks are limited to two or three GRNs on which all inference tools exhibit very low accuracy levels, due in no small part to the incompleteness of those GRNs. On the other hand, there has been a surge in development of realistic simulators of expression data, including those that simulate the dynamics of a GRN using biophysical principles and incorporate realistic models of expression noise [13, 37]. This presents the opportunity of utilizing simulated data sets and their underlying GRNs as training data for supervised GRN reconstruction. This is the key idea explored in our work.

Motivated by the above considerations, we develop here a new GRN inference tool called SPREd, which trains a neural network model to predict GRN edges from expression profiles. It relies on a very large training set comprising millions of TF-gene pairs, generated using a state-of-the-art expression simulator. Systematic evaluations show that SPREd outperforms leading methods such as GENIE3, ENNET, TIGRESS and PORTIA on unseen synthetic data sets. We also find that SPREd generalizes well when tested on synthetic data with different characteristics from those used in training. Finally, we demonstrate that SPREd is more accurate than other methods on the most reliable and widely used “real data” benchmark available today. We believe SPREd is a promising first step towards more advanced implementations of supervised GRN reconstruction where our rapidly developing understanding of GRN dynamics are encoded in increasingly realistic simulators, which then lead to even more powerful training sets for training highly expressive ML models.

## RESULTS

### SPREd: supervised learning framework for GRN

The GRN inference problem addressed here is to detect the TFs regulating a given gene based on expression data across a range of “conditions”. (We will use “conditions” to refer to different experimental conditions, replicates, cell types or cells.) Leading methods use ML for this task, modeling the expression of the target gene as a function of the expression levels of all TFs, then examining the contribution of each TF (feature) to the model and reporting the important TFs as regulators of the gene (**Figure 1A** top). In other words, current methods approach GRN prediction via a related problem – gene expression prediction – and extract the GRN as a “by-product” of solving that problem. In contrast, we use a more direct, supervised learning approach to GRN prediction (**Figure 1A** bottom). Instead of relating TFs’ expression (input) to target gene expression (output), we relate the TFs’ *and* target gene’s expression (input) to TF-gene edges (output). Learning such a relationship requires training data comprising many input-output pairs, each of which comprises an expression matrix (input) and its corresponding GRN (output). Since reliable GRNs are rarely available, we used a simulator that constructs a realistic but synthetic expression matrix corresponding to a GRN, repeating the process many times, and used the resulting expression matrix-GRN pairs to train the GRN prediction model. Once such a model has been trained, it can be presented expression data that has not been seen before and directly predict the GRN that those data may have arisen from.

**Figure 1.**
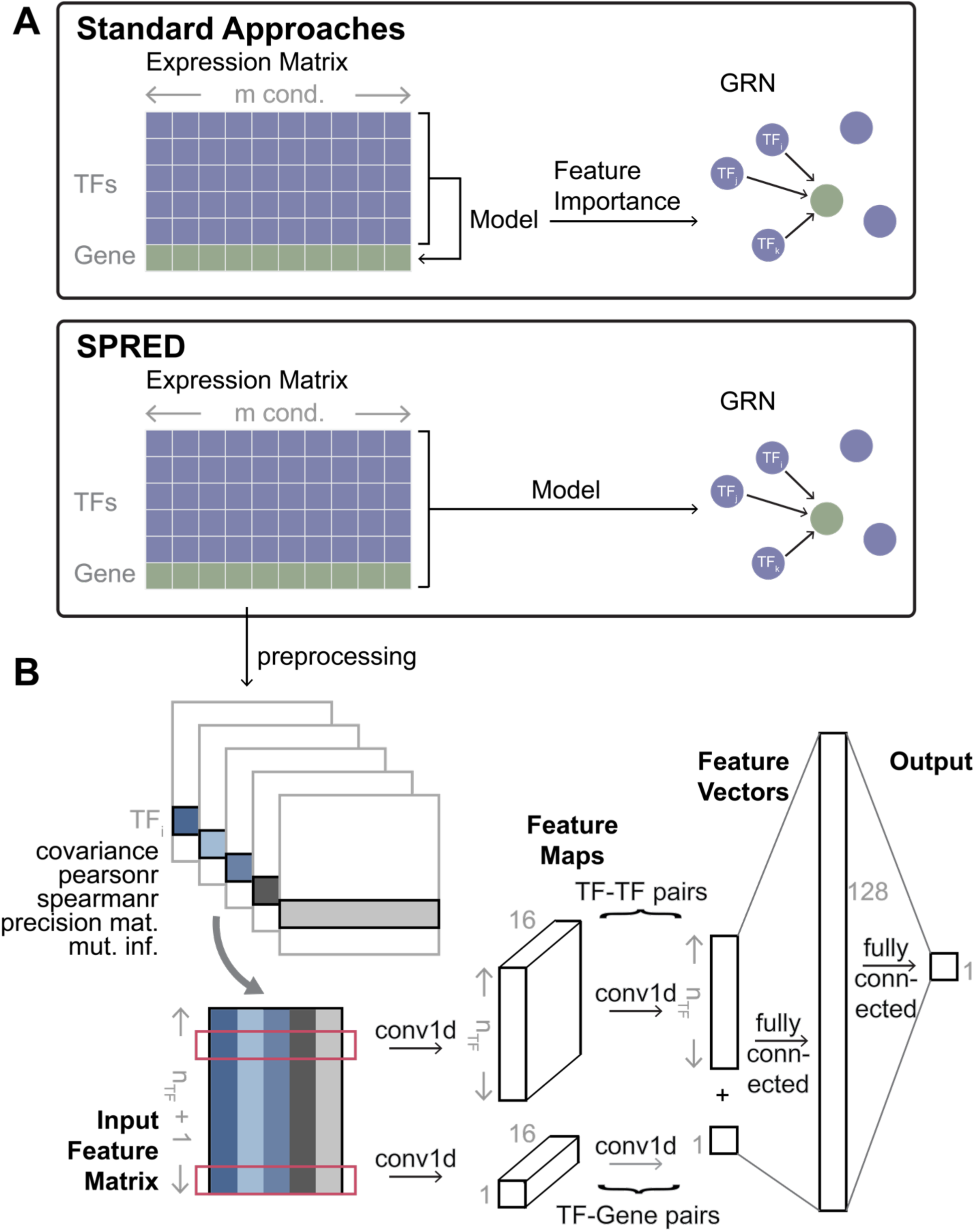
SPRED: a simulation-supervised learning framework for gene regulatory network (GRN) inference. **(A)** Standard approaches typically build ML models of the target genes using the expression levels of TFs as features. GRNs are then constructed based on the feature importance of TFs (features) in the model trained for a target gene. In SPREd, an ML model is trained to directly predict TFs regulating a target gene, based on expression matrix of all TFs and the target gene. The ML model is trained on simulated expression matrix-GRN pairs and can then be used to predict the GRN for any expression matrix. **(B)** Architecture of SPRED-SP neural network model. Given an expression matrix whose rows represent *n_TF_* TFs and one target gene (panel A), the preprocessing step creates five matrices of pairwise relations (features) for each TF-TF pair and each TF-target gene pair. These features include covariance, Pearson correlation, Spearman correlation, mutual information and precision matrix entry corresponding to the TF-TF or TF-target gene pair. The five features of every gene pair involving a particular TF (say TF_i_) then serve as the inputs of a 1D convolutional neural network (CNN) (input feature matrix, size (*n_TF_* + 1) × 5). The feature map resulting from the first layer of convolution (out channel = 16) is of dimension (*n_TF_* + 1) × 16, and feeds into a second convolution layer, whose outputs are fully connected to a hidden layer, which finally connect to the output layer. The output layer consists of a binary label indicating if TF_i_ is a regulator of the target gene (details shown in Methods).

Our method, called SPREd (<u>S</u>upervised <u>P</u>redictor of <u>R</u>egulatory <u>Ed</u>ges), utilizes a neural network to relate an expression matrix to the corresponding GRN. Here, we will assume that the expression matrix includes the expression levels of a single target gene and all the candidate TFs across a set of *m* conditions, and the GRN consists of a subset of the *n_TF_* candidate TFs that are regulators of the target gene (**Figure 1A**). We implemented two complementary models, “SPREd-MultiLabel” (SPREd-ML) and “SPREd-SinglePair” (SPREd-SP), for the GRN prediction task. These two models have similar architectures and feature definitions but differ in one key aspect: SPREd-SP predicts regulatory relationships one TF-gene pair at a time, while SPREd-ML simultaneously scores all *n_TF_* candidate TFs for their regulatory relationship to a gene. For both models, the input expression matrix is represented via features that capture the relationships between TF(s)-target gene pairs and TF-TF pairs. For SPREd-SP, when predicting the relationship between *TF_i_* and gene *g*, features are calculated for the pair (*i*, *g*) and (*i*, *j*) for every candidate *TF_j_*, yielding a total of *n_pairs_* = 1 + *n_TF_*. For SPREd-ML, since the output of the model includes relationships between every candidate TF and gene *g*, features are calculated for all *n_TF_*

TFs paired with gene *g* and for all 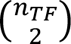 TF-TF pairs, yielding 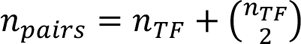. For each pair, five statistics including covariance, Pearson correlation, Spearman correlation, mutual information, and “precision score” are calculated (see Methods). These five statistics for each of the TF-gene and TF-TF pairs are presented as a *n_pairs_* × 5 matrix of “features” in the input layer of a 1d convolutional neural network (CNN). The architecture of the neural network (**Figure 1B** and **Supplementary Figure S8**), with convolution layer(s) producing a *n_pairs_*-dimensional vector (convolving the five features of each pair into one value), followed by fully connected hidden layers to the output layer, which consists of one unit for SPREd-SP or *n_TF_* units for SPREd-ML with values of 0-1 indicating the absence or presence of an edge between the TF and the target gene.

The SPREd neural network is trained on synthetic data generated using the SERGIO simulator. SERGIO (**Figure 2A**) takes an input GRN, with a subset of TFs marked as master regulators (MRs) that are TFs without input regulators thereof. Starting with the expression levels of the MRs, SERGIO simulates the dynamics of transcriptional regulation and samples cells from the steady state distributions of those dynamics. By varying the MRs’ expression profiles, we can diversify the biological conditions simulated by SERGIO. To generate training data for the SPREd model, we created GRNs with a special structure (**Figure 2A, left**). Each GRN has a set of *n_G_* target genes, regulated by a set of *n_TF_* TFs, which are in turn regulated by *n_MR_* MRs. A gene has *d*_*TF*→*g*_ incoming regulatory edges, while each TF is under the control of *d*_*MR*→*TF*_ MRs. This simple 3-layer structure allows us to control the complexity of combinatorial regulation and global co-expression in the simulated expression data used for model training and testing. To generate an expression matrix of *m* conditions, we used SERGIO to simulate *m* cells specified by *m* distinct MR profile (see Methods). This produces an expression matrix with *n_MR_* + *n_TF_* + *n_G_* rows and *m* columns, which is then processed to generate training samples for SPREd (see Methods), with *n_TF_n_G_* samples being created for SPREd-SP and *n_G_* samples for SPREd-ML. The whole process is repeated *n_GFN_* times, each time creating a GRN at random with the desired connectivity properties.

**Figure 2.**
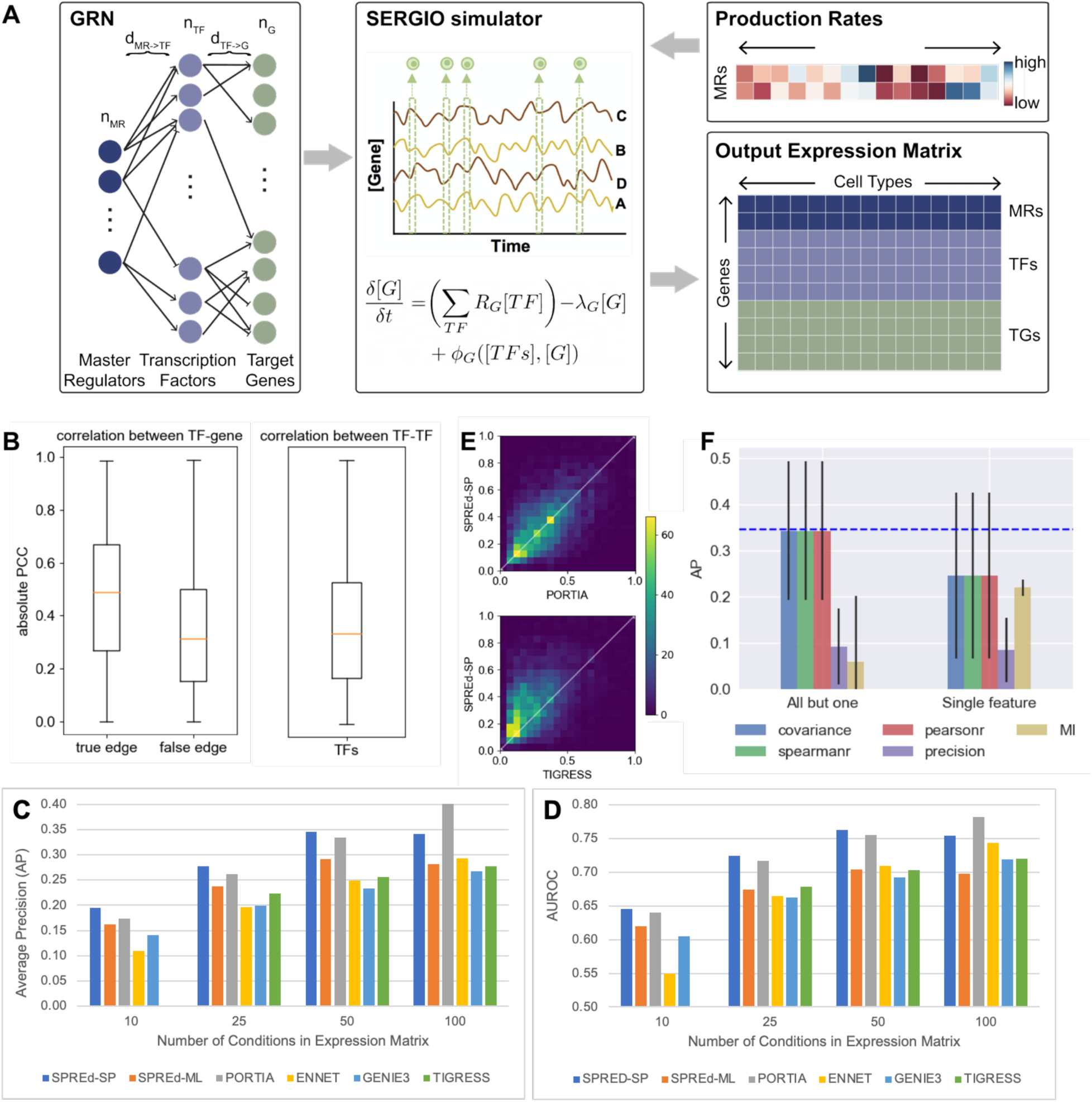
SPRED exhibits superior performance on synthetic datasets. **(A)** Schematic of synthetic data generation. Each synthetic data set comprises a GRN (left) and an expression matrix (bottom right). The GRN has three “layers” – master regulators (MR), transcription factors (TF) and target genes, with regulatory edges from one layer to the next. The MRs are included as the first layer so as to induce co-expression among TFs, mimicking real data. Parameters describing the GRN include the number of MRs (*n_MR_*), the number of transcription factors (*n_TF_*), the number of target genes (*n_G_*), the number of incoming edges to each target gene (*d*_TF→G_), and the number of incoming edges to each TF (*d*_MR→TF_). A GRN is sampled at random while respecting these parameters and is used by SERGIO (middle), a biophysics-based model, to simulate the expression profiles of different artificial biological conditions, each of which is described by the production rates of MRs (top right), thus generating an expression matrix whose rows include target genes, TFs and MRs and columns represent biological conditions. **(B)** Co-expression statistics of synthetic expression data. Absolute value of Pearson correlation coefficient (PCC) of TF-gene pairs (left) that comprise GRN edges (“true edges”) and those that do not (“false edges”) and of TF-TF pairs (right). Results are from simulations using GRNs in default configuration. **(C,D)** Average precision or “AP” (**C**) and AUROC (**D**) of the six evaluated methods – SPREd-SP, SPREd-ML, PORTIA, ENNET, GENIE3 and TIGRESS – on data sets with varying numbers of conditions (columns in expression matrix). Each performance metric (AP or AUROC) is calculated for individual target genes, and results summarized over 5000 genes from 50 GRNs (100 genes in each GRN). **(E)** Direct comparison of average precision (AP) between SPRED and PORTIA (top) or TIGRESS (bottom), for expression data with 50 conditions. **(F)** AP of SPREd-SP when using all but one (left) or only one (right) of the five features describing each TF-gene or TF-TF pair. AP when using all five features is shown by blue dashed line.

### SPREd outperforms leading GRN inference methods on synthetic benchmarks

We first evaluated the prediction accuracy of SPREd on a synthetic benchmark that resembles the training data described above. This benchmark comprises 50 GRNs, each with *n_G_* = 100 target genes, and thus presents 50 × 100 = 5000 “test samples”, each being a GRN with a single target gene and its known regulator TFs, and the corresponding expression matrix to be used as input. A GRN reconstruction method must predict, based on the expression matrix for each test sample, which of the *n_TF_* candidate regulators actually regulate the target gene. In general, such methods assign scores to each candidate regulator and their accuracy can be assessed using standard metrics such as Area Under Receiver Operating Characteristic (AUROC) or Area Under Precision Recall Curve (AUPRC). Here, in addition to the AUROC, we relied on the Average Precision (AP, see Methods) score rather than the AUPRC, because it is better suited for extremely sparse positive sets. As comparators for evaluating predictive accuracy, we relied on four state-of-the-art methods – ENNET [27], GENIE3 [25], TIGRESS [22] and PORTIA [38], which have emerged as top methods in recent benchmarking studies [38] and represent a diversity of modeling approaches.

In the first benchmark, we set the number of conditions *m* to 50 and the GRN parameters to *n_MR_* = 5, *d*_*MR*→*TF*_ = 3, *n_TF_* = 100 and *d*_*TF*→*G*_ in the range [3,7]. The number of conditions is typical for bulk transcriptomics data from an individual PI’s laboratory (a few experimental conditions and 3-5 replicates per condition [39]). It is also the ballpark number of samples (individuals) for each tissue in GTEx [40]. The parameters *n_MR_* and *d*_*MR*→*TF*_ determine the level of co-expression among TFs, an important characteristic of the benchmark since extensive co-expression among TFs in real data sets is one of the major hurdles in accurate GRN prediction. **Figure 2B** shows the extent of TF-TF correlation in the generated benchmark, which is comparable to that of TF-gene pairs that are not GRN edges; GRN edges have significantly greater correlations on average, as expected, but we also note a substantial overlap between the distributions of edges and non-edges, pointing to the difficulty of the GRN prediction task for this benchmark. We trained SPREd on a data set of *n_GFN_* = 250 GRNs generated with the same simulation parameters as the test set. As shown in **Figure 2C** (group “50”) and **Table 1A** (*m* = 50), the AP of SPREd-SP is higher on average than those of all other methods, with strong statistical significance in most cases (**Supplementary Table S1A**). Similarly, SPREd-SP has a higher AUROC than these methods except for PORTIA, to which it is comparable (**Figure 2D, Table 1B, Supplementary Table S1B**). **Figure 2E** shows head-to-head comparisons of SPREd-SP with the two next-best competing methods (PORTIA and TIGRESS), revealing higher AP score of SPREd-SP versus TIGRESS for the majority of test samples, and a noticeable degree of complementarity between SPREd-SP and PORTIA. We noted SPREd-ML to have weaker performance than SPREd-SP and PORTIA in these evaluations, though it exhibited higher AP and AUROC than ENNET, GENIE3 and TIGRESS.

**Table 1.** Performance comparison of SPREd, PORTIA, ENNET, GENIE3 and TIGRESS. Average Precision (AP) (**A**) and AUROC (**B**) in evaluations on expression matrices with varying number of conditions (columns in expression matrix). Each performance metric (AP or AUROC) is calculated for individual target genes, and results averaged over 5000 genes from 50 GRNs (100 genes in each GRN). Expression data are simulated using GRNs with *n_MR_*=5, *n_TF_*=100, *d*_*TF*→*G*_ = 3-7. Note: TIGRESS runs for 10 conditions did not complete successfully.

| #conditions | SPREd-SP | SPREd-ML | PORTIA | ENNET | GENIE3 | TIGRESS |
| --- | --- | --- | --- | --- | --- | --- |
| 10 | **0.19** | 0.16 | 0.17 | 0.11 | 0.14 | - |
| 25 | **0.28** | 0.24 | 0.26 | 0.20 | 0.20 | 0.22 |
| 50 | **0.35** | 0.29 | 0.33 | 0.25 | 0.23 | 0.26 |
| 100 | 0.34 | 0.28 | **0.40** | 0.29 | 0.27 | 0.28 |

**Table 1. Performance comparison of SPREd, PORTIA, ENNET, GENIE3 and TIGRESS.**
| #conditions | SPREd-SP | SPREd-ML | PORTIA | ENNET | GENIE3 | TIGRESS |
| --- | --- | --- | --- | --- | --- | --- |
| 10 | **0.65** | 0.62 | 0.64 | 0.55 | 0.61 | - |
| 25 | **0.72** | 0.67 | 0.72 | 0.67 | 0.66 | 0.68 |
| 50 | **0.76** | 0.70 | 0.76 | 0.71 | 0.69 | 0.70 |
| 100 | 0.75 | 0.70 | **0.78** | 0.74 | 0.72 | 0.72 |

We next repeated the above evaluations at additional values of *m* (number of conditions) and observed the performance advantage of SPREd-SP to persist when there are fewer conditions (*m* = 25, 10) (**Table 1, Figure 2C,D, Supplementary Table S1**), though with *m* = 100, the performance of PORTIA improves over SPREd-SP. Similarly, the above-noted advantage of SPREd-ML over ENNET, GENIE3 and TIGRESS persists on data with fewer conditions but disappears at *m* = 100. (Also see **Supplementary Figure S1**). This is in line with our expectation that with sufficient sample size to train models of gene expression, the need for an alternative paradigm such as the synthetic data-supervised approach of SPREd diminishes. In this regime, the ability to tune free parameters on the given expression matrix can give an edge to the standard approach (Figure 1A) in vogue today. We note that where this regime begins (e.g., ∼100 conditions in our benchmark) depends on the data set and has to be estimated with care.

We also observed similar relative performances when comparing SPREd-SP with other tools on the DREAM5 benchmark of [36], see Supplementary Table S2, S3. In particular, we observed SPREd performance to be the best among compared methods in terms of AP, for three of the four networks, the exception being the network with the largest ratio of number of conditions (samples) to TFs (covariates).

We next determined the importance of the five features representing each gene pair to the accuracy of SPREd-SP. Firstly, we repeated the evaluations using only one of the features and found that the performance is substantially below that of the full model for any one of the features (**Figure 2F**, “Single feature”). This suggested that the model utilizes complementary information from multiple features. We also evaluated subsets comprising four of the features, leaving out one at a time, and observed a substantial drop only when leaving out precision (**Figure 2F**, “All but one”), indicating that the information carried by each of the remaining features is largely redundant with at least one other feature (e.g., Pearson correlation and Spearman correlation are expected to be largely mutually redundant), whereas the precision score likely captures information complementary to the others. (See **Supplementary Figure S2** for similar analysis with SPREd-ML.)

### Effect of benchmark diversity on predictive accuracy

We next examined how SPREd performance varies with benchmark generation parameters that determine the difficulty of the GRN inference task. First, we looked at the effect of edge density between TFs and genes (*d*_*TF*→*G*_), set to 3 − 7 above. We repeated the evaluations with the range changed to [1,2] (less combinatorial regulation) or to [8,10] (more combinatorial). One might expect that as the edge density increases, the AP score should increase: if a gene has many regulators, then even a randomly predicted regulator is likely to be correct. However, we observed (**Figure 3A**) that the AP is highest in the *d*_*TF*→*G*_ ∈ [1,2] (least combinatorial) setting. (Also see **Supplementary Figures S3, S4**.) One possible reason for this is that with greater numbers of true regulators of a gene, mutual correlations among TFs (**Figure 2B**) exacerbate the multicollinearity problem, making the TF prediction task harder. The literature suggests that the more complex cases of combinatorial transcriptional regulation (e.g., in early development) involve 3 − 7 TFs regulating a gene [41], but the rare examples of comprehensively reconstructed GRNs (e.g., in yeast) suggest an edge density of 1 − 2 on average. These observations provide some real biological context to the evaluations above.

**Figure 3.**
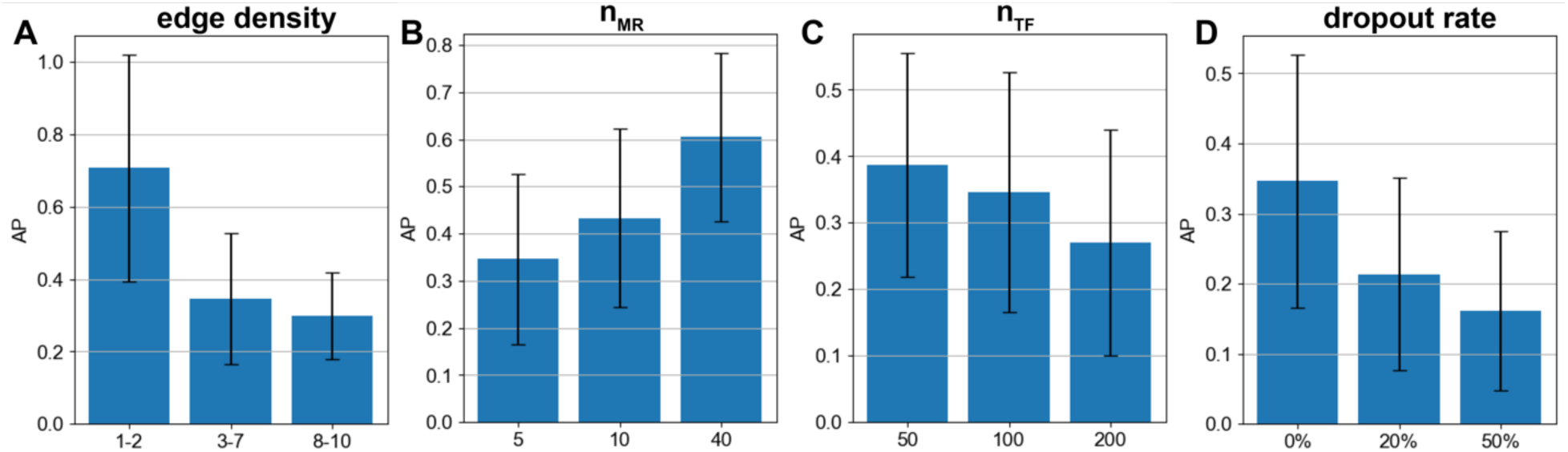
Effect of benchmark parameters on GRN reconstruction. Performance (average precision) of SPREd-SP with varying edge density (dTF→G) of 1-2, 3-7, and 8-10 TFs per target gene (**A**), varying numbers of MRs (nMR) (**B**), varying numbers of TFs (nTF) (**C**), and varying levels of dropout added to the synthetic expression matrix (**D**).

Recall that our motivation in adding the “master regulator” layer in synthetic GRNs is that it provides an easy way to control the extent of TF-TF correlations, a major challenge for GRN recovery. (In a more realistic, “cascade-like” GRN, two TFs’ mutual correlation will depend on their relative position in the cascade.) In our 3-layer GRNs, as *n_MR_* increases, the pairwise correlation of TFs decreases and we found the GRN reconstruction to become easier, as expected (**Figure 3B**) .

The parameter *n_TF_* directly controls the difficulty of the GRN reconstruction task, since the task involves selecting the true regulators among all *n_TF_* candidate TFs. Clearly, if we increase *n_TF_* while keeping *d*_*TF*→*G*_ fixed, then for each gene the number of positives to be determined remains the same but the number of candidates increases, so the AP will decrease, which is what we observed (**Figure 3C**). We noted however less than the expected 2-fold decrease in AP when we double the number of TFs, from 50 to 100 and then from 100 to 200. This suggests increasing *n_TF_*leads to an unexplained improvement in ability to identify true regulators, countered by a stronger deterioration due to the larger pool of candidate TFs.

Finally, we examined the effect of expression measurement noise beyond the stochasticity of gene expression already modeled in the SERGIO simulation process. In particular, we simulated a simplistic version of “dropout” [42] whereby a certain fraction of entries of the expression matrix are artificially at zero values. As expected, the performance deteriorates with increasing noise (**Figure 3D**) and at a dropout level of 50% the performance of SPREd-SP drops to nearly random expectation (**Supplementary Figure S4**). This corroborates previous reports that dropouts, unless properly mitigated by imputation tools [43–45], can be a major problem for GRN reconstruction [37].

In the above evaluations, the two SPREd models, unlike the other four methods, are supervised with training data that has the same characteristics as the test data, which might give them an advantage. To partially offset this advantage, we repeated the comparisons on more heterogeneous benchmarks, with GRNs in the test set being created with a mixture of all settings of *n_MR_* or *d*_*TF*→*G*_ parameters. We found (**Figure 4B**) that the advantages of SPREd models over comparators, seen above for mid-range edge densities (**Table 1**), are as strong or greater with the broader spectrum of edge densities. Similarly, when benchmarking with a broader span of *n_MR_* values (5, 10, 40) that induce varying levels of TF-TF correlations, the advantage of SPREd over PORTIA and GENIE3 persists (**Figure 4A**), though the gap between SPREd and ENNET/TIGRESS shrinks. We repeated these evaluations with SPREd trained on GRNs with the benchmark parameter (*n_MR_* or *d*_*TF*→*G*_) set to a specific value while the test set remained a heterogeneous mix of GRNs, and found the model’s performance to remain unchanged (**Supplementary Figures S5A,B**). These results indicate that the performance of the model depends primarily on the difficulty of the test set rather than the precise characteristics of the GRNs used in training, which is a sign of generalizability.

**Figure 4.**
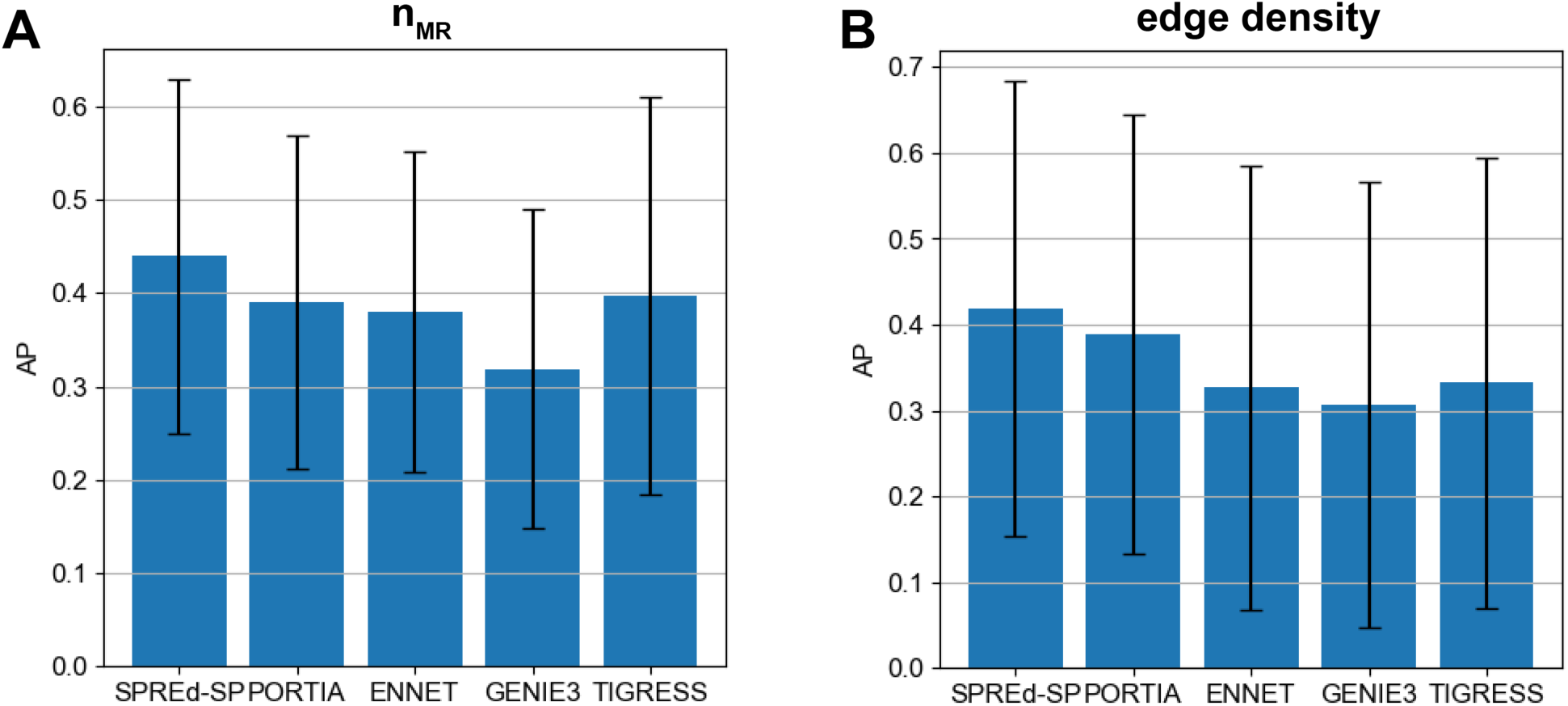
Performance comparison on heterogeneous benchmarks. AP comparison of SPRED-SP, SPRED-ML, PORTIA, ENNET, GENIE3 and TIGRESS on heterogeneous datasets comprising GRNs with **(A)** *n_MR_* =5, 10 and 40 (in equal numbers) and **(B)** with *d*_*TF*→*G*_ = 1, 2, … 10 (in equal numbers).

### SPREd performance is better than or competitive with leading methods on real data

Finally, we performed comparative evaluations of the six methods on one of the best characterized eukaryotic GRNs available today, from the yeast *S. cerevisiae*, reconstructed based on multiple types of cis-regulatory information, and compiled in the form of benchmarks by Siahpirani & Roy [46]. The Siahpirani-Roy yeast benchmark includes three “gold standard GRNs” named “MACISAAC2” [47], “YEASTRACT TYPE2” AND “YEASTRACT COUNT3” (see Methods), based on cis-regulatory evidence including TF ChIP-seq profiles, TF motif matches and evolution conservation and have since been used by other studies [38]. The benchmark also includes three different expression matrices, named “NatVar”, “Knockout”, and “StressResp”, reflecting expression variation under different axes of biological differences. We evaluated the six methods for each of the nine combinations of the expression matrix as input and the gold standard GRN as ground truth, called “tests” below. Each test involves identifying typically 1 − 2 TFs regulating a target gene (out of ∼250 candidate TFs), for ∼2,000 target genes.

First, we examined AP values (**Table 2A**) and noted that the SPREd models have the highest median performance across the nine tests, with either one yielding the highest AP on six of the tests and PORTIA exhibiting the greatest AP on the remaining three. A head-to-head comparison (**Supplementary Figure S6**) reveals that both models clearly perform better than or similar to ENNET, GENIE3 and TIGRESS across the tests, while both models are competitive with PORTIA with benchmark-dependent complementarity. Examination of AUROC values (**Table 2B**) revealed that the two SPREd models have the highest median performance across tests, with SPREd-ML being the best among all methods for seven of the tests and SPREd-SP and GENIE3 being the best in one test each. **Supplementary Figure S7** reveals the clear advantage of SPREd-ML over all other methods, while SPREd-SP outperforms ENNET and TIGRESS and is competitive with PORTIA and GENIE3.

**Table 2.**
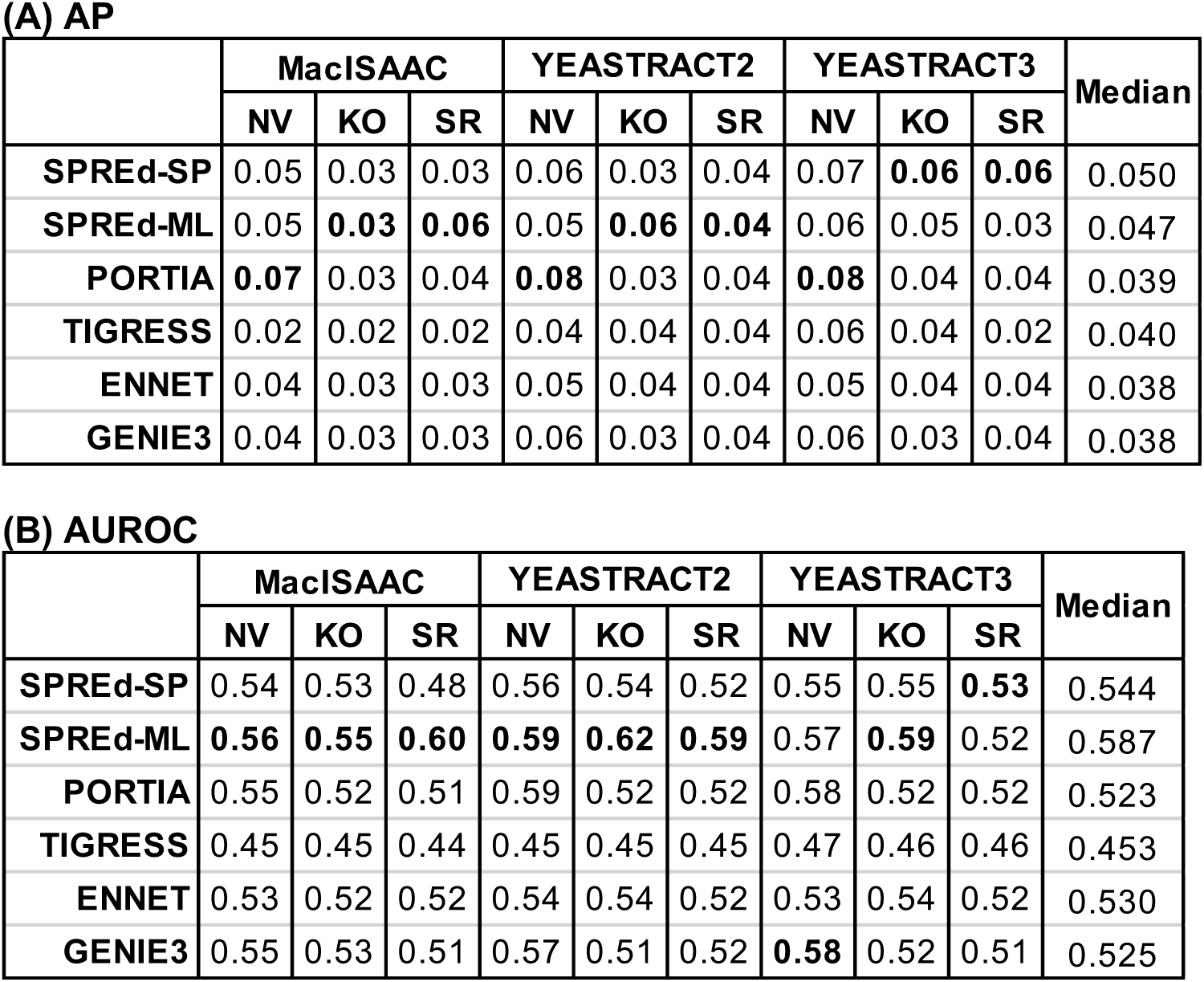
Mean AP (**A**) and AUROC (**B**) of six evaluated methods on benchmarks using the MacIsaac2, YEASTRACT type2 (YEASTRACT2), or YEASTRACT count3 (YEASTRACT3) gold standard networks and the Nat Var (NV), Knockout (KO), and StressResp (SR) expression data from Siahpirani & Roy. AP/AUROC values are calculated for individual genes and averaged over genes. Highest value in each column is shown in bold. Median value across the nine benchmarks is shown for each method in the rightmost column.

**(A) AP**
|  | MacISAAC |  |  | YEATRACT2 |  |  | YEATRACT3 |  |  | Median |
| --- | --- | --- | --- | --- | --- | --- | --- | --- | --- | --- |
|  | NV | KO | SR | NV | KO | SR | NV | KO | SR |  |
| **SPREd-SP** | 0.05 | 0.03 | 0.03 | 0.06 | 0.03 | 0.04 | 0.07 | **0.06** | **0.06** | 0.050 |
| **SPREd-ML** | 0.05 | **0.03** | **0.06** | 0.05 | **0.06** | **0.04** | 0.06 | 0.05 | 0.03 | 0.047 |
| **PORTIA** | **0.07** | 0.03 | 0.04 | **0.08** | 0.03 | 0.04 | **0.08** | 0.04 | 0.04 | 0.039 |
| **TIGRESS** | 0.02 | 0.02 | 0.02 | 0.04 | 0.04 | 0.04 | 0.06 | 0.04 | 0.02 | 0.040 |
| **ENNET** | 0.04 | 0.03 | 0.03 | 0.05 | 0.04 | 0.04 | 0.05 | 0.04 | 0.04 | 0.038 |
| **GENIE3** | 0.04 | 0.03 | 0.03 | 0.06 | 0.03 | 0.04 | 0.06 | 0.03 | 0.04 | 0.038 |

**Table 2.** Mean AP (**A**) and AUROC (**B**) of six evaluated methods on benchmarks using the MacIsaac2, YEASTRACT type2 (YEASTRACT2), or YEASTRACT count3 (YEASTRACT3) gold standard networks and the Nat Var (NV), Knockout (KO), and StressResp (SR) expression data from Siahpirani & Roy.
|  | MacISAAC |  |  | YEATRACT2 |  |  | YEATRACT3 |  |  | Median |
| --- | --- | --- | --- | --- | --- | --- | --- | --- | --- | --- |
|  | NV | KO | SR | NV | KO | SR | NV | KO | SR |  |
| **SPREd-SP** | 0.54 | 0.53 | 0.48 | 0.56 | 0.54 | 0.52 | 0.55 | 0.55 | **0.53** | 0.544 |
| **SPREd-ML** | **0.56** | **0.55** | **0.60** | **0.59** | **0.62** | **0.59** | 0.57 | **0.59** | 0.52 | 0.587 |
| **PORTIA** | 0.55 | 0.52 | 0.51 | 0.59 | 0.52 | 0.52 | 0.58 | 0.52 | 0.52 | 0.523 |
| **TIGRESS** | 0.45 | 0.45 | 0.44 | 0.45 | 0.45 | 0.45 | 0.47 | 0.46 | 0.46 | 0.453 |
| **ENNET** | 0.53 | 0.52 | 0.52 | 0.54 | 0.54 | 0.52 | 0.53 | 0.54 | 0.52 | 0.530 |
| **GENIE3** | 0.55 | 0.53 | 0.51 | 0.57 | 0.51 | 0.52 | **0.58** | 0.52 | 0.51 | 0.525 |

We noted that the AP values in Table 2A are relatively low (in the 5% range) in absolute terms, regardless of the method. This is consistent with previous studies [38] and is likely due to the sheer difficulty of the problem where only a handful of ∼250 candidate regulators are known to be positives. Indeed, when we reexamined the AP values in light of the random baseline, we found up to 2.4-fold higher than expected AP values for SPREd predictions (**Supplementary Table S6**). For example, in the three tests noted where SPREd-ML achieves an AP of ∼0.06, this value is 2.1-fold, 2.3-fold and 2.4-fold greater than random. At the same time, the relatively low fold changes (in AP) over random expectation in several of the other tests suggests that even the best methods available today leave ample scope for improvement. It is also likely that the true regulators of a typical gene are largely missing from the gold standard GRNs and the precision values are underrepresented as a result.

## DISCUSSION

SPREd in its current form is designed and tested for GRN inference from bulk RNA-seq data, which, despite the spreading popularity of single cell technologies, remains the platform of choice when a number of samples in varying biological conditions have to be profiled, especially on average laboratory budgets but also in consortium efforts [40]. As such, our performance comparisons have been focused on state-of-the-art tools for the bulk data domain, which form an actively growing genre in its own right. Despite obvious parallels, GRN inference for bulk and single-cell transcriptomics data presents distinct challenges – far smaller sample sizes in the bulk case and far greater technical noise in the single-cell case. They also address two different biological notions of GRN: single-cell GRNs explain gene expression variation from cell to cell [48], which may in part be due to cell type differences, while bulk-based GRNs explain expression variation across biological conditions or individuals. An obvious task cut out for the future is to explore the SPREd approach for single-cell GRN inference; such efforts may begin with training models on synthetic single-cell data generated using SERGIO in ‘noisy’ mode but will likely need to explore alternative model architectures as well.

Several avenues of future improvements to SPREd present themselves. The current model uses as features five different types of pairwise relationships between genes, three of which (covariance, Pearson correlation, Precision matrix) capture linear relationships, one (Spearman correlation) captures monotonic relationships and one (mutual information) is more general. It will be interesting to explore if a 2D convolution of the joint distribution between two genes, akin to the approach of [29], can improve the featurization of pairwise relationships used by SPREd’s neural network. Finally, given that SPREd does not perform parameter training, which is a strategy successfully used by its competing methods, a combination of these two complementary approaches will be a promising direction for future research.

GRN reconstruction from expression data is a very challenging problem, in part because the goal is one of causal TF-gene relationships based on observational data alone, and also because the structure of real GRNs leads to extensive correlations among the TFs themselves. To overcome these fundamental obstacles, other kinds of evidence such as cis-regulatory elements or TF knockout/knockdown data are used in influential studies [46, 49–51], but such data may not always be easily available, which necessitates continued attention to the core problem of expression-to-GRN mapping. A second major challenge facing GRN reconstruction is the scarcity of reliable “gold standard” GRNs corresponding to the available expression data. A GRN edge (TF-gene pair), to be considered true, should at the very least be supported by evidence of gene dysregulation upon perturbing the TF’s expression and by credible evidence of functional TF binding to an enhancer linked with that gene. To our knowledge, there are very few, if any, GRNs that meet these stringent criteria of credibility *and* are reasonably complete (i.e., many or most edges known). This is the reason why synthetic benchmarks remain the primary evaluation approach today [37, 52, 53]. At the same time, it is expected that method developers evaluate their GRN reconstruction tools on real data sets, but one should not be surprised if such evaluations reveal low accuracies, given the shortcomings of current GRN benchmarks in terms of completeness and/or soundness. Indeed, we and others [38, 46] have consistently noted relatively low values of GRN reconstruction accuracy on real data benchmarks.

Our comparative evaluation involved four methods that have been found in a recent benchmarking study as leading methods for the task [38] and employ a variety of techniques (Random Forests, Gradient Boosting, Least Angle Regression, Precision Matrix calculation). Notably, only one of these tools (PORTIA) [38] is recent, while the others are over 10 years old. This reflects the reality that recent related work has not focused on the core problem of GRN inference from expression data, instead aiming to exploit additional types of data (e.g., epigenomic profiles and motifs) or tackling the noisy nature of single-cell data.

The performance of GRN inference methods generally does not depend on the number of target genes in the network, but it does depend on the number of candidate TFs and the number of actual regulators of a gene. The largest number of TFs considered in any of our tests is about 330 TFs, which is much smaller than the ∼2000 TFs in human. However, in practice one expects GRN inference tools to be run with a selected subset of TFs, e.g., those with detectable expression or differential expression in the samples of interest. We recommend such pre-selection as it may help improve the accuracy of inferred GRN edges, and caution against using SPREd with thousands of TFs as candidate regulators.

Several avenues of future improvements to SPREd present themselves. Firstly, systematic tuning of hyperparameters of the neural network model should allow us to train more accurate models, especially if combined with even larger numbers of training samples, which are easy to generate. Secondly, we note that the current model uses as features five different types of pairwise relationships between genes, three of which (covariance, Pearson correlation, Precision matrix) capture linear relationships, one (Spearman correlation) captures monotonic relationships and one (mutual information) is more general. Our ablation analysis (Supplementary Figure S2) suggested that these five measures have mutual redundancies and it may be possible to remove one or more of these features. We leave this as a future modification, to be evaluated extensively in varying benchmark settings. Moreover, it will be interesting to explore if a 2D convolution of the joint distribution between two genes, akin to the approach of [29], can improve the featurization of pairwise relationships used by SPREd’s neural network. We used a CNN architecture in SPREd as it is better able to suitably combine the five different measures of pairwise expression relationships between TF-TF pairs and TF-gene pairs, compared to a simpler MultiLayer Perceptron model (Supplementary Table S5); however, future work may explore alternative and more effective architectures. Finally, given that SPREd does not perform parameter training, which is a strategy successfully used by its competing methods, a combination of these two complementary approaches will be a promising direction for future research.

GRN inference methods in the literature have generally been evaluated using a common benchmark called “DREAM5 GRN challenge” [36]. We chose the 3-layer GRNs and corresponding SERGIO-simulated expression datasets as our primary benchmark as we wanted our evaluations to systematically vary different parameters that control the difficulty of the problem. Note that the expression data simulator (SERGIO in “clean” mode) we used to generate the benchmark is very similar to the GeneNetWeaver [54] simulator used in DREAM. The main difference between the DREAM benchmarks and ours is in the underlying GRNs, which are far more numerous and varied in our benchmark.

## METHODS

### Simulation of expression matrix from a GRN, using SERGIO

The SERGIO simulator [37] requires “targets file” describing the regulators of each gene in the GRN and the “master regulators file” with information on MRs as inputs. The targets file lists the non-MR gene’s identifier, the number of regulators of that gene, their identifiers, interaction parameter (*K*) representing the strength and directionality of influence of corresponding regulators, and Hill coefficients representing the degree of non-linearity of the influence (see SERGIO documentation). Each *K* was sampled from a uniform distribution on [1.0, 5.0], and then negated with 0.2 probability. Each Hill coefficient was set to 2 with 0.9 probability and 1 with 0.1 probability. The number of “cell types” to be simulated was set to 100, and the number of cells per cell type was set to 1 . The master regulators file requires specification of each MR’s production rate in each cell type. This was done by first selecting, for each GRN, a pair of ranges defining low and high rates respectively, and then sampling each MR’s rate in each cell type at uniform from these ranges. (The pair of ranges were as suggested by SERGIO and chosen uniformly at random.) Given the targets file and master regulators file, the SERGIO simulator produces steady-state expression matrices with dimension of (*n_MR_* + *n_TF_* + *n_G_*) × 100. Each of the *m* “cell types” simulated by SERGIO was treated as a separate “condition”. In benchmarks where *m* < 100, *m* of the 100 simulated “cells” were selected at random as conditions.

### Performance evaluation metrics

All evaluations were performed for each target gene separately and averaged over genes. For a gene, the task is to predict whether each of the *n_TF_* candidate regulators is actually a regulator, i.e., a binary classification task on *n_TF_* test samples, allowing computation of AUROC. Instead of AUPRC, we used the related metric “Average precision” [55] *AP* = ∑*_n_*(*R*_*n*_ − *R*_*n−1*_)*P_n_*, where *R_n_* and *P_n_* denote the *n^th^* threshold of recall and precision. GRNs in benchmarks used here have very few regulators per gene, which may lead to misleading AUPRC values, especially in the extreme but common case of one true regulator.

### Input data preparation in SPREd

Given an expression matrix with *n_TF_* rows representing TFs and one row for the target gene, and *m* columns representing conditions, the first step is to use a Box-Cox power transformation [56] on each gene, followed by z-transformation resulting in zero mean and unit standard deviation for each gene. (In case the expression matrix has any negative values, all entries are shifted by a constant to achieve strictly positive values.) Pairs of rows of the resulting normalized expression matrix were examined to calculate a covariance matrix, denoted as Σ, Pearson correlation coefficient matrix, Spearman correlation coefficient matrix, and discrete mutual information matrix, each of dimensionality *N* × *N*, where *N* = *n_TF_* + 1. The *N* × *N* precision matrix was calculated using (Σ + ε × *I*)^−1^, ε = 1e − 3 . Since the resulting matrices are symmetric, 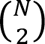 + *N* distinct entries of each matrix are used as features for the SPREd-ML model. The SPREd-SP model, on the other hand, predicts only whether the *i^th^* TF is a regulator of the target gene or not, and its features include row *i* of the above five matrices. For both models, all pairwise relationship features of each of the five types are flattened into a vector.

### Architecture of SPREd-ML

The SPREd-ML architecture is shown in **Supplementary Figure S8**. The input of the neural network is a *n_pairs_* × 5 matrix as described above. The *n_pairs_* rows of this matrix include 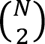 rows representing each TF-TF pair and *N* rows representing each TF-target gene pair, the target gene being fixed. Each of the former set of 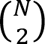 rows feeds into a shared 1D convolutional kernel, while each of the latter set of *N* rows feeds into another shared 1D convolutional kernel. These two kernels result in feature maps of dimensions 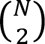 × 1 and *N* × 1 respectively, which are fully connected to *n_TF_* units of the output layer. We used rectified linear activation (ReLU) as the activation function throughout the network. Connections between the feature map representing TF-target gene pairs and the output layer had a dropout rate of 0.3.

### Architecture of SPREd-SP

The overall architecture is presented in **Figure 1B**. The input is a *n_pairs_* × 5 matrix, with the *n_pairs_* = *N* = *n_TF_* + 1 rows including *n_TF_* rows for TF-TF pairs (one of the TFs being fixed) and one row for the TF-target gene pair. These inputs feed into two 1D convolution-batchnorm-ReLU blocks with dimensions 5 × 16 and 16 × 1 respectively. (The *n_TF_* rows for TF-TF pairs and the single row for the TF-gene pair are convolved using separate kernels.) The resulting feature map is a *n_pairs_*-dimensional vector that is fully connected to a 128-node layer, which in turn is fully connected with the single output node.

### SPREd training details

Both models are trained using Adam optimizer with weight decay of 5e-4 and learning rate of 2e-4. Binary cross entropy with logits is used as the loss function. The positive weight is set to 9, the batch size is set to 32 and the maximum number of training epochs is set to 300. The training was performed on an Nvidia V100 GPU. See Supplementary Table S4 for samples of run-times of training and application of trained models (testing) of SPREd.

### Availability and implementation

Data and code are available from https://github.com/iiiime/SPREd.

### Key points

- SPREd is a tool for gene regulatory network (GRN) inference from expression data.
- The underlying model is a convolutional neural network that directly predicts regulators of a gene.
- The model is trained on large data generated by a state-of-the-art GRN simulator, and no model training is performed on the user’s data.
- SPREd outperforms competing methods on comprehensive benchmarks involving synthetic as well as real datasets.

## ACKNOWLEDGEMENTS

This work was supported by the National Institutes of Health [R35GM131819A to S.S.].

