## supplementary figures and tables for "SPREd: A simulation-supervised neural network tool for gene regulatory network reconstruction"

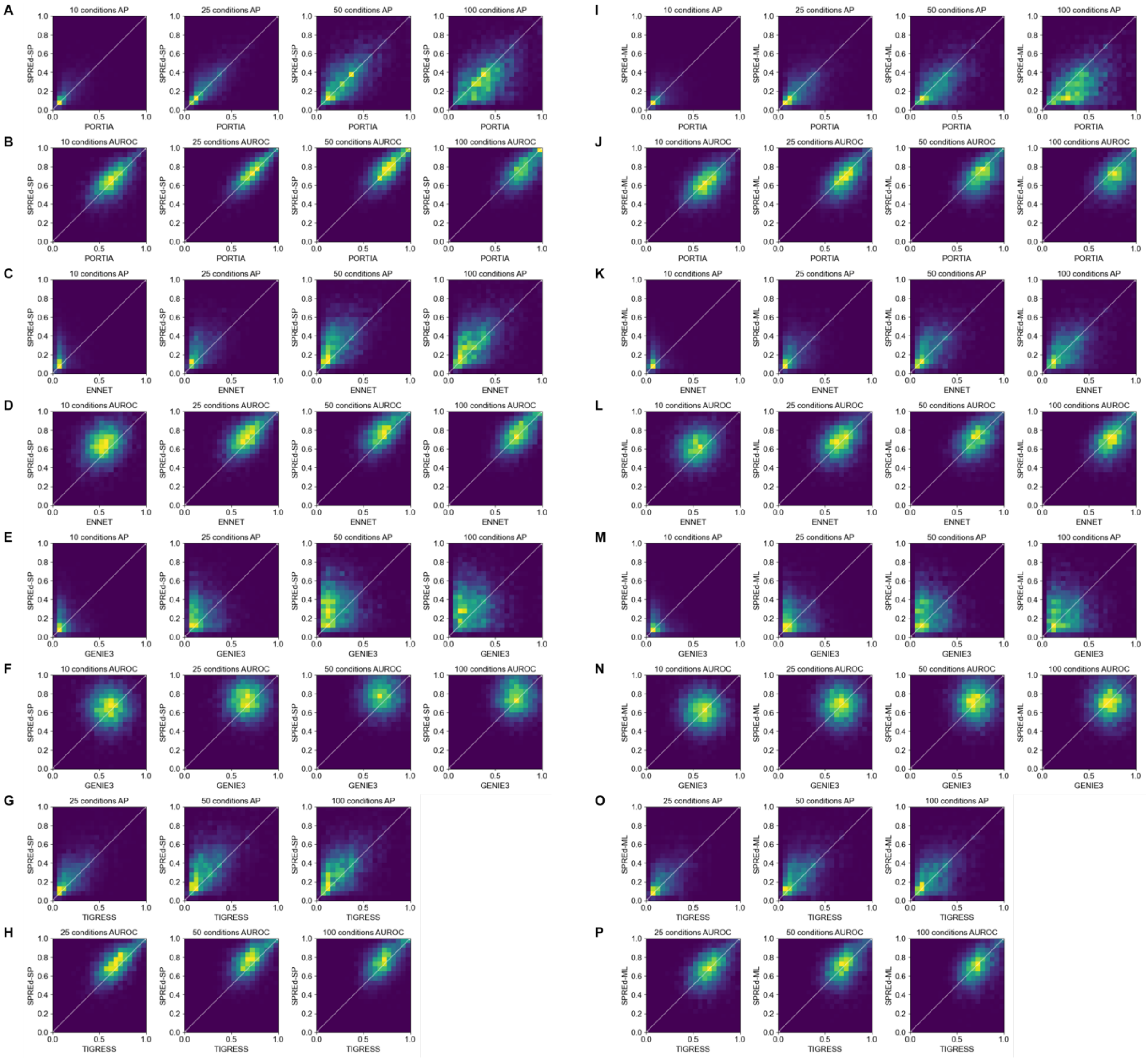


**Supplementary figure S1.** Direct comparisons of AP (**A**, **C**, **E**, **G**, **I**, **K**, **M**, **O**) and AUROC (**B**, **D**, **F**, **H**, **J**, **L**, **N**, **P**) between SPREd-SP and PORTIA (**A**, **B**), SPREd-SP and and ENNET (**C**, **D**), SPREd-SP and GENIE3 (**E**, **F**), SPREd-SP and TIGRESS (**G**, **H**), SPREd-ML and PORTIA (**I**, **J**), SPREd-ML and ENNET (**K**, **L**), SPREd-ML and GENIE3, or between SPREd-ML and TIGRESS (**O**, **P**).

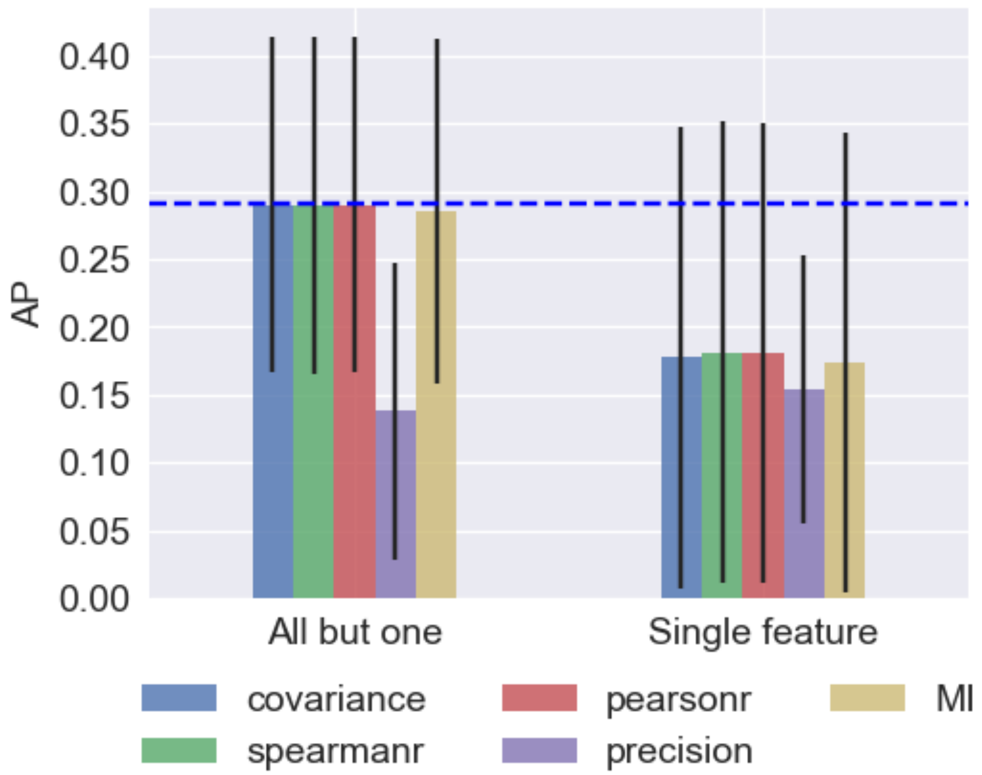


**Supplementary figure S2.** Average Precision of SPREd-ML when using all but one (left) or only one (right) of the five features describing each TF-gene or TF-TF pair. AP when using all five features is shown in blue dashed line.

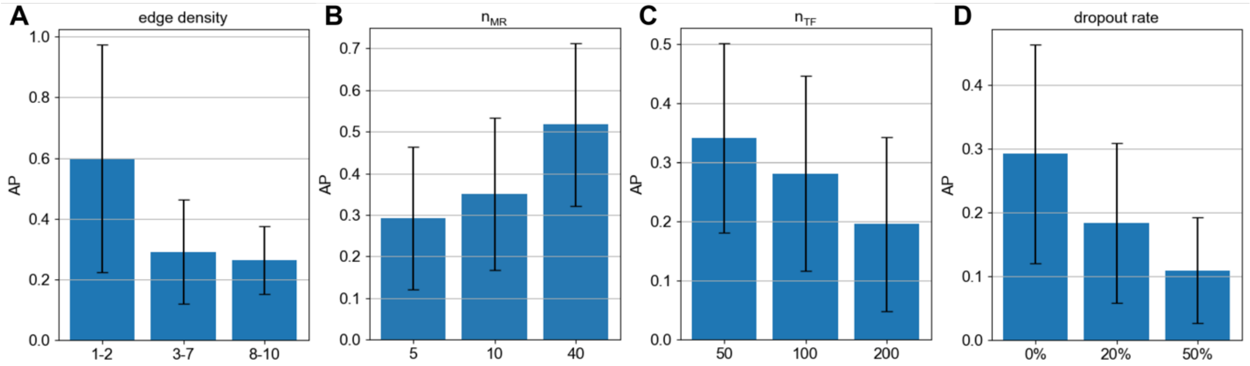


**Supplementary figure S3. AP scores of SPREd-ML for varying benchmark parameters.** Performance (average precision) of SPREd-ML with varying edge density ($d_{TF\rightarrow G}$) of 1-2, 3-7, and 8-10 TFs per target gene (**A**), varying numbers of MRs ($n_{MR}$) (**B**), varying numbers of TFs ($n_{TF}$) (**C**), and varying levels of dropout added to the synthetic expression matrix (**D**).

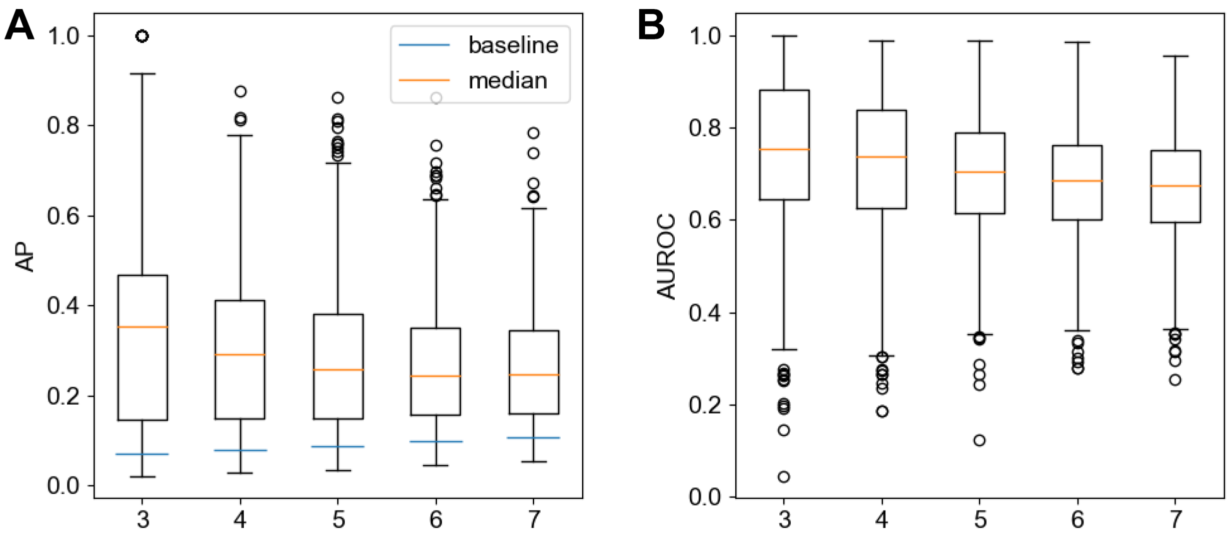


**Supplementary figure S4.** SPREd performance (with default GRN parameters of $n_{MR} = 5$, $n_{TF} = 100$, $d_{MR\rightarrow TF}$ = 2) at different values of edge density $d_{TF\rightarrow G}$ = 3, 4, … 7. Average Precision (AP) is shown in panel (A) and AUROC is shown in panel (B). The blue line in (A) shows the random expectation of AP.

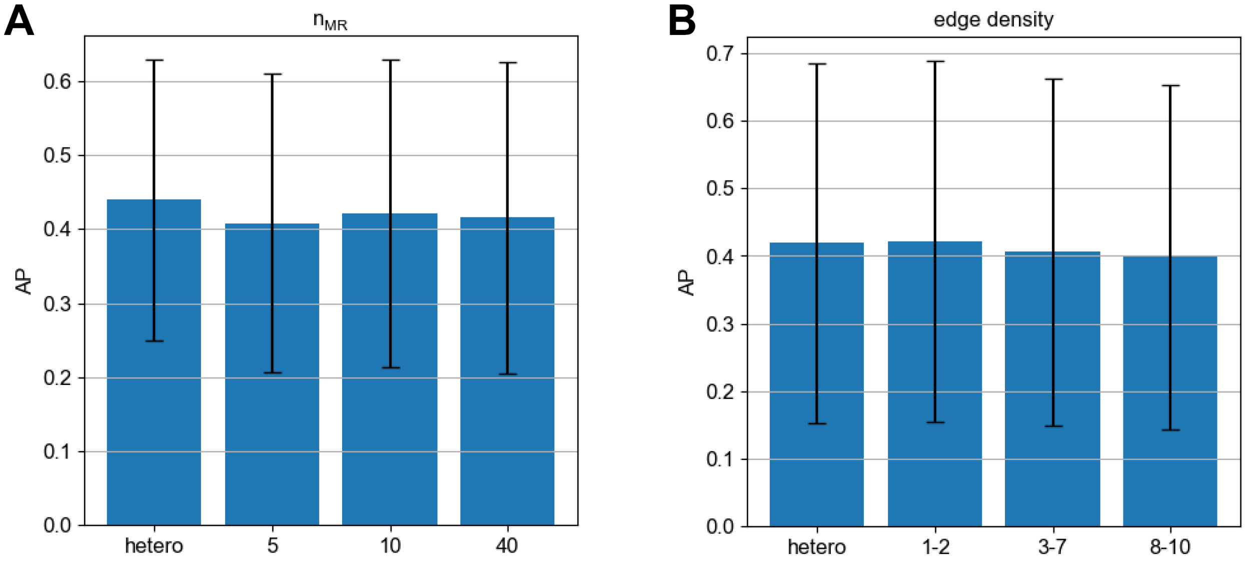


**Supplementary figure S5. Performance comparison on heterogeneous benchmarks.** (**A**) Performance of SPREd-SP on heterogeneous test sets of different using model weights from training sets with $n_{MR}$ set to 5, 10, and 40 or a mix thereof ("hetero"). (**B**) Performance of SPREd-SP on heterogeneous test sets of varying edge densities using model weights trained on datasets with different edge densities – 1-2, 3-7, 8-10, or the entire range ("hetero").

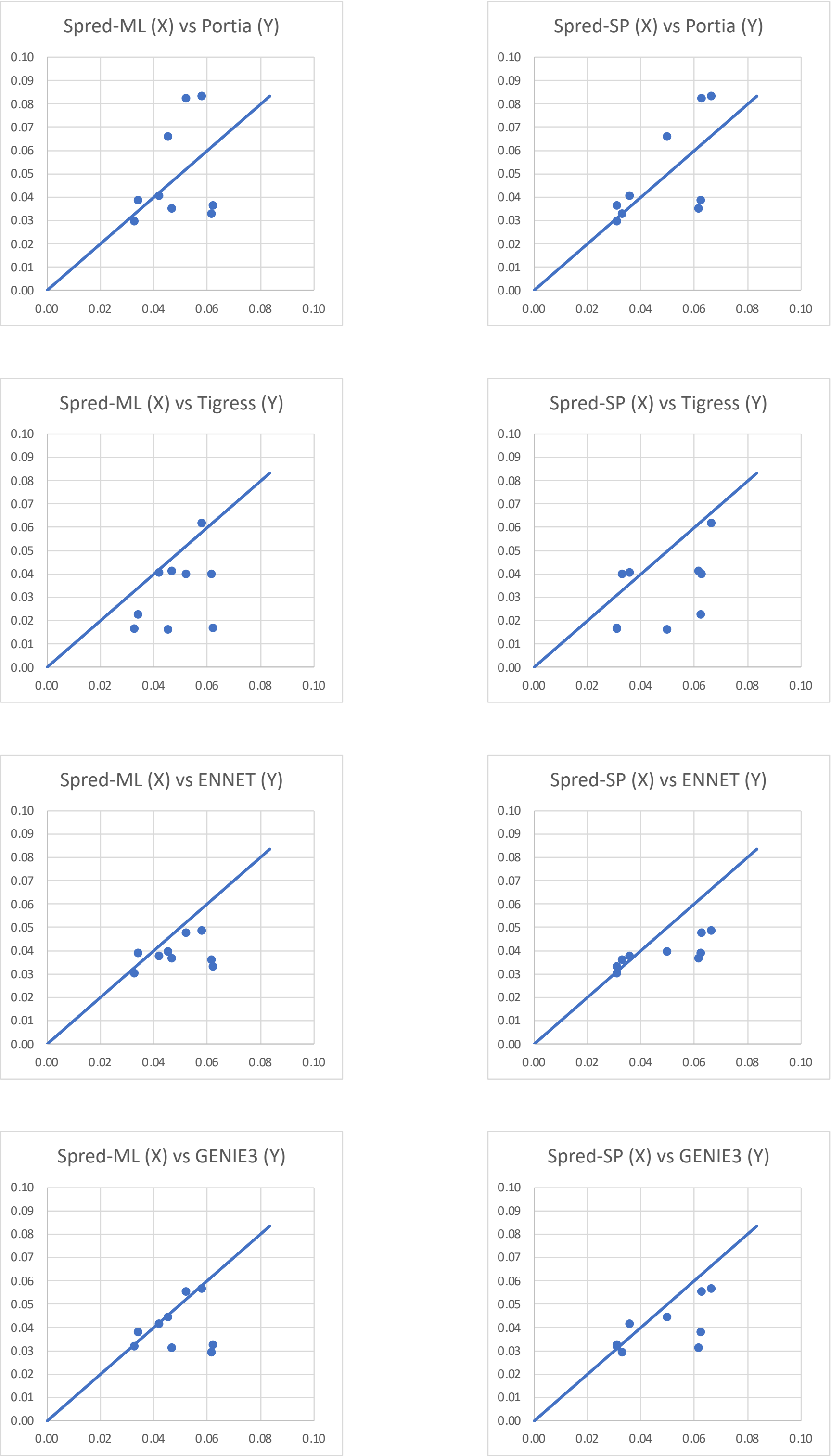


**Supplementary figure S6.** Head-to-head comparison of SPREd-ML (left) or SPREd-SP (right) versus each comparator (PORTIA, TIGRESS, ENNET, GENIE3) in terms of Average Precision (AP) on each of the nine benchmarks ("tests"). In each panel, the nine points shown represent the tests, X-axis is the AP (averaged over all genes) of a SPREd model and Y-axis is the AP of a comparator.

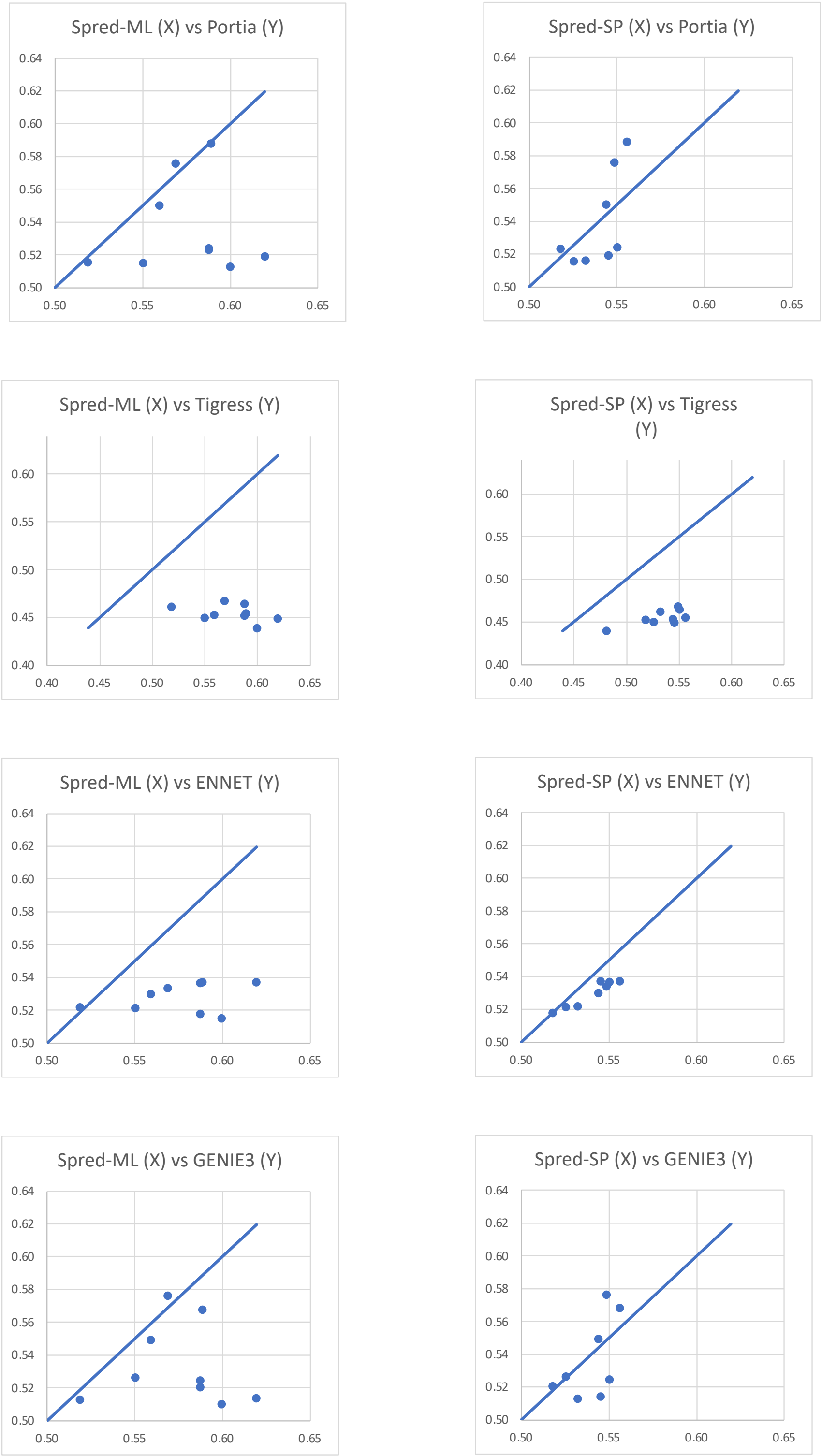


**Supplementary figure S7.** Head-to-head comparison of SPREd-ML (left) or SPREd-SP (right) versus each comparator (PORTIA, TIGRESS, ENNET, GENIE3) in terms of AUROC on each of the nine benchmarks (“tests”). In each panel, the nine points shown represent the benchmarks, X-axis is the AUROC (averaged over all genes) of a SPREd model and Y-axis is the AUROC of a comparator.

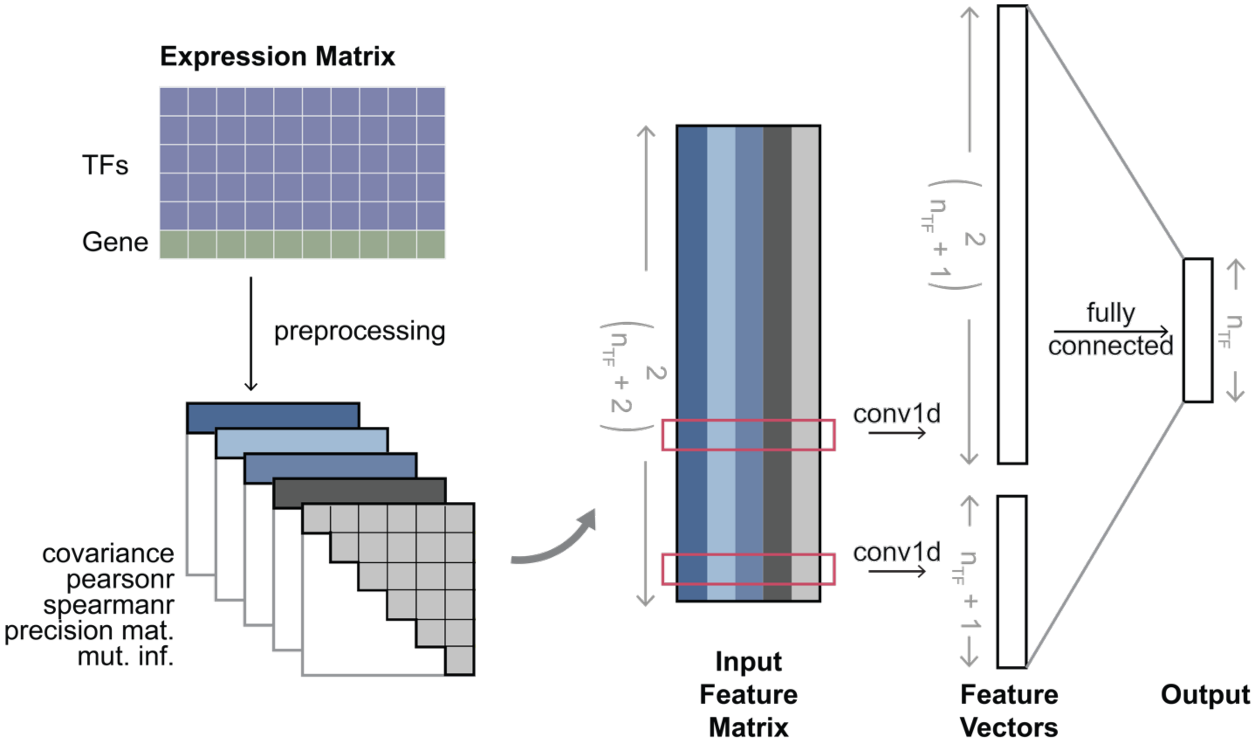


**Supplementary figure S8. Architecture of SPREd-ML neural network model.** Given an expression matrix whose rows represent $n_{TF}$ TFs and one target gene, the preprocessing step creates five features for each of the $\binom{n_{TF}+1}{2}$ TF-TF pairs and each of the $n_{TF}+1$ TF-target gene pairs. These features include covariance, Pearson correlation, Spearman correlation, mutual information and precision matrix entry corresponding to the TF-TF or TF-target gene pair. The five features of every gene pair then serve as the inputs of a 1D convolutional neural network (CNN). The output layer consists of $n_{TF}$ binary labels indicating if each TF is a regulator of the target gene.

(**A**)

| | PORTIA | ENNET | GENIE3 | TIGRESS |
|---|---|---|---|---|
| **100** | 6.74E-63 | 6.29E-45 | 3.68E-106 | 2.42E-71 |
| **50** | 0.0024 | 4.34E-179 | 1.15E-245 | 2.74E-144 |
| **25** | 2.01E-07 | 1.92E-175 | 5.13E-161 | 1.33E-68 |
| **10** | 8.89E-15 | 0 | 9.90E-108 | - |

(**B**)

| | PORTIA | ENNET | GENIE3 | TIGRESS |
|---|---|---|---|---|
| **100** | 1.09E-26 | 1.07E-05 | 4.98E-44 | 4.28E-43 |
| **50** | 0.00043 | 8.32E-97 | 3.29E-160 | 9.16E-123 |
| **25** | 0.00055 | 9.97E-108 | 1.28E-117 | 8.00E-65 |
| **10** | 0.0024 | 3.95E-250 | 1.02E-51 | - |

**Supplementary table S1.** P-values of paired Wilcoxon test (two-tailed) of difference in AP scores (**A**) and AUROC (**B**) on 5000 genes between SPREd-SP and PORTIA, ENNET, GENIE3, or TIGRESS. Green/red font indicates better/worse SPREd-SP performance.

| | $n_{TF}$ | $\mathbf{n_{genes}}$ | $\mathbf{n_{cond}}$ | $\mathbf{n_{targets}}$ |
|---|---|---|---|---|
| **Network1** | 195 | 1643 | 805 | 1387 |
| **Network2** | 99 | 2810 | 160 | 367 |
| **Network3** | 334 | 4511 | 805 | 922 |
| **Network4** | 333 | 5950 | 536 | 1798 |

**Supplementary table S2.** Summaries of different networks in the DREAM5 benchmark, including the number of TFs ($n_{TF}$), the number of total genes ($n_{genes}$), the number of conditions ($n_{cond}$), and the number of target genes ($n_{targets}$) for each network. Source: https://www.synapse.org/#!Synapse:syn2787209/wiki/70350

(**A**)

| | SPREd-SP | PORTIA | GENIE3 | ENNET |
|---|---|---|---|---|
| **Network1** | 0.31 | 0.52 | **0.58** | 0.035 |
| **Network2** | **0.16** | 0.12 | 0.13 | 0.055 |
| **Network3** | **0.20** | 0.17 | 0.16 | 0.017 |
| **Network4** | **0.053** | 0.039 | 0.034 | 0.017 |

(**B**)

| | SPREd-SP | PORTIA | GENIE3 | ENNET |
|---|---|---|---|---|
| **Network1** | 0.71 | 0.85 | **0.87** | 0.50 |
| **Network2** | 0.61 | 0.61 | **0.63** | 0.53 |
| **Network3** | 0.67 | 0.67 | **0.71** | 0.49 |
| **Network4** | 0.54 | 0.54 | **0.55** | 0.50 |

**Supplementary table S3.** Performance evaluation on different networks from the DREAM5 benchmark. We compared the performance of different methods in terms of Average Precision (AP) (**A**) and AUROC (**B**). TIGRESS was not included here since it has a runtime issue and the runs failed to complete.

**(A) Runtime for training**

| $n_{TF} \times n_{genes}$ | **Time (s) per epoch** |
|---|---|
| 2500000 | 378.45 |
| 1500000 | 222.87 |
| 500000 | 76.741 |

**(B) Runtime for testing**

| $n_{TF} \times n_{genes}$ | **Time (s)** |
|---|---|
| 500000 | 558.43 |
| 100000 | 109.45 |
| 50000 | 54.52 |

**Supplementary table S4.** Run-time and scalability of SPREd-SP. (**A**) Run time for training the SPREd model, per epoch, for varying data sizes ($n_{TF} \times n_{genes}$). A full training typically spans 50 epochs. (**B**) Run time for applying a trained SPREd model to infer GRN for data sets of varying sizes.

| | AP | AUROC |
|---|---|---|
| **SPREd** | 0.35 | 0.76 |
| **MLP** | 0.26 | 0.70 |
| **p-value** | 0 | 1.23e-310 |

**Supplementary table S5**. Comparison of AP score and AUROC between SPREd-SP CNN architecture and simple 2-layer MLP. Pair-wise Wilcoxon p-values comparing SPREd and MLP model are listed at the bottom.

| | MacIsaac2 | Yeastract Type2 | Yeastract Count3 |
|---|---|---|---|
| **NatVar** | 1.54 | 1.74 | 2.14 |
| **KO** | 1.19 | 2.28 | 1.66 |
| **Stress** | 2.39 | 1.54 | 1.25 |

**Supplementary table S6.** Mean of "fold change AP" of SPREd-ML over all genes, for each of the nine benchmarks. Fold change AP refers to the observed AP divided by the random baseline AP for a gene.
